# HILPDA uncouples lipid storage in adipose tissue macrophages from inflammation and metabolic dysregulation

**DOI:** 10.1101/566802

**Authors:** Xanthe A.M.H. van Dierendonck, Montserrat A. de la Rosa Rodriguez, Anastasia Georgiadi, Frits Mattijssen, Wieneke Dijk, Michel van Weeghel, Rajat Singh, Jan Willem Borst, Rinke Stienstra, Sander Kersten

**Author notes:** Correspondence to: Nutrition, Metabolism and Genomics group Division of Human Nutrition Wageningen University Stippeneng 46708 WE Wageningen The Netherlands E-mail or. These authors contributed equally to this work (Xanthe A.M.H. van Dierendonck, Montserrat A. de la Rosa Rodriguez). Shared senior authors (Rinke Stienstra, Sander Kersten).

## Abstract

Obesity promotes accumulation of lipid-laden macrophages in adipose tissue. Here, we determined the role of macrophage lipid accumulation in the development of obesity-induced adipose tissue inflammation, using mice with myeloid-specific deficiency of the lipid-inducible HILPDA protein. HILPDA deficiency in bone marrow-derived macrophages markedly reduced intracellular lipid levels and accumulation of fluorescently-labeled fatty acids in lipid droplets. Decreased lipid storage in HILPDA-deficient macrophages could be almost completely rescued by inhibition of adipose triglyceride lipase (ATGL) and was associated with increased oxidative metabolism. In diet-induced obese mice, HILPDA deficiency did not alter inflammatory or metabolic parameters, despite markedly reducing lipid storage in adipose tissue macrophages. Our data indicate that HILPDA is a lipid-induced physiological inhibitor of ATGL-mediated lipolysis that uncouples lipid storage in adipose tissue macrophages from inflammation and metabolic dysregulation. Overall, our data question the importance of lipid storage in adipose tissue macrophages in obesity-induced inflammation and metabolic dysregulation.

## INTRODUCTION

The occurrence of obesity among the world population has risen tremendously over the past 50 years and has become a huge public health concern. Obesity is characterized by excess adipose tissue mass and is associated with a state of chronic low-grade inflammation in several metabolic tissues, including adipose tissue. This low-grade inflammation has been suggested to be an important pathophysiological mechanism underlying many of the adverse health effects associated with obesity^1^.

The main role of adipose tissue is to serve as a depot for surplus energy via the storage of lipids. The homeostasis of adipose tissue is crucial for maintaining whole body insulin sensitivity. During the development of obesity, adipose tissue homeostasis is disturbed, altering the production of several inflammatory cytokines and adipokines. It is believed that the increase in inflammatory cytokines and adipokines can disrupt normal insulin signalling and contribute to obesity-associated insulin resistance^2^.

Macrophages are key innate immune cells that are important for maintaining homeostasis in healthy adipose tissue, but also contribute to the development of inflammation during obesity^3^. In lean states, adipose tissue macrophages predominantly show anti-inflammatory phenotypes and are distributed evenly throughout the adipose tissue. In contrast, in obese adipose tissue, macrophages accumulate in so-called crown-like structures around dead adipocytes and display a metabolically activated phenotype^4–6^. Metabolically activated macrophages residing in these crown-like structures form multiple intracellular lipid droplets and display distinct transcriptional profiles involving lysosomal lipolysis^6–8^. This feature is indicative of an attempt by macrophages to buffer excess lipids, which is adaptive in the lean state but becomes maladaptive in obese adipose tissue^3^. The presence of foam cell-like macrophages in obese adipose tissue is reminiscent of lipid-laden macrophages described in the context of atherosclerotic plaques. Although foam cell formation and adipose tissue inflammation are known to co-exist during obesity, the exact role of lipid accumulation in adipose tissue macrophages in the development of obesity-induced adipose tissue inflammation and associated metabolic disturbances remains unclear.

HILPDA is a small lipid droplet-associated protein that is expressed in several tissues (Mattijssen, 2014). The expression of *Hilpda* is induced by a number of different stimuli including hypoxia, beta-adrenergic activation, and PPARs. Gain and loss of function studies have shown that HILPDA promotes lipid deposition in hepatocytes, adipocytes and macrophages^9–13^. The mechanism by which HILPDA promotes lipid storage in cells has not been completely elucidated but evidence has been presented that HILPDA directly binds and inhibits adipose triglyceride lipase (ATGL)^14^, consistent with the ability of HILPDA to inhibit lipolysis^10,11^. Interestingly, endothelial cell marker *Tie2*-Cre driven deletion of *Hilpda* was found to decrease fatty acid and oxLDL-driven lipid droplet formation in macrophages, and reduce lesion formation and progression of atherosclerosis in ApoE-/- mice^13^.

Inasmuch as HILPDA increases intracellular lipid accumulation, modulation of its expression will aid in deciphering the role of lipid accumulation in adipose tissue macrophages in the development of obesity-induced adipose tissue inflammation. Here, we aimed to determine the exact role of HILPDA in lipid accumulation in macrophages, and explore the potential causal relationship between lipid accumulation in adipose tissue macrophages and the development of adipose tissue inflammation and insulin resistance during obesity.

## RESULTS

### Hilpda as gene of interest in obese adipose tissue

To identify genes that may be able to modify lipid storage in adipose tissue macrophages, we searched for lipid-induced genes that are both induced by lipids and elevated in adipose tissue macrophages upon obesity. To that end, we co-analyzed the transcriptomics data from the following three experiments: 1) adipose tissue macrophages isolated from obese mice versus lean mice, 2) mouse peritoneal macrophages treated with the fatty acid oleate, 3) mouse peritoneal macrophages treated with intralipid, a triglyceride emulsion. Scatter plot analysis led to the identification of *Hilpda* as a gene of particular interest (Fig 1A), as it was the only gene that was strongly upregulated in all three experiments. Moreover, *Hilpda* was the most highly induced gene in peritoneal macrophages by oleate treatment. Consistent with a potential role for *Hilpda* in obese-induced adipose tissue inflammation, transcriptomics analysis indicated that the expression of *Hilpda* in adipose tissue is upregulated during high fat feeding in mice in parallel with macrophage and inflammatory marker genes, such as *Ccl2* (MCP1), *Cd68*, and *Itgax* (Cd11c) (Fig 1B), which was confirmed by qPCR (Fig 1C). Immunohistochemistry of adipose tissue of obese mice indicated that HILPDA co-localized with adipose tissue macrophages found in crown-like structures, thus supporting the production of HILPDA by adipose tissue macrophages (Fig 1D). Together, these data suggest that HILPDA may be implicated in obesity-induced adipose tissue inflammation and foam cell formation.

**Figure 1.**
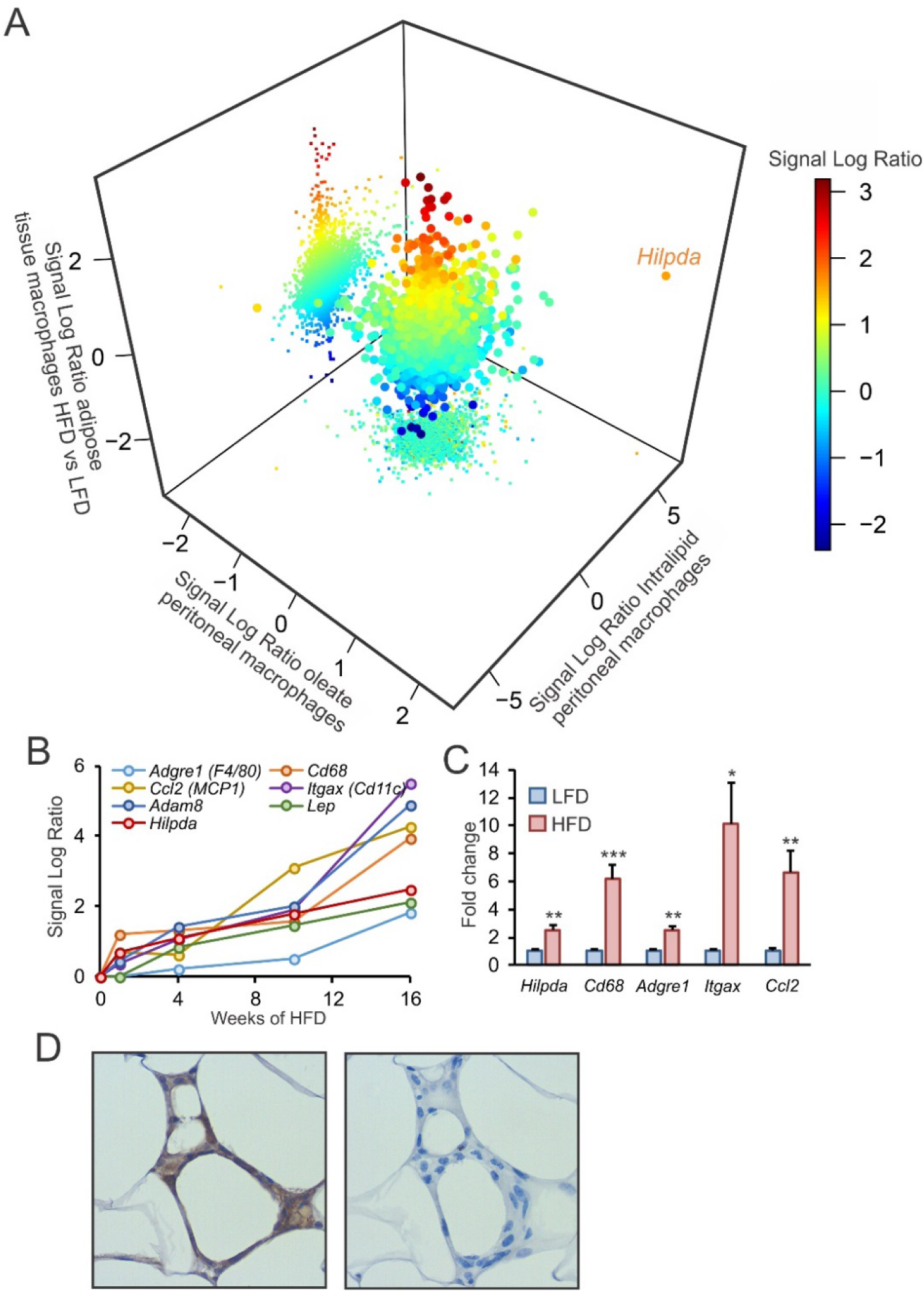
*Hilpda* as gene of interest in obese adipose tissue. A: Transcriptomics co-analysis of ATMs isolated from C57Bl/6 mice fed a HFD versus a LFD, C57Bl/6 mouse peritoneal macrophages treated with oleate and C57Bl/6 mouse peritoneal macrophages treated with intralipid. B: Signal log ratio patterns of *Hilpda*, inflammatory genes and macrophage markers in gonadal adipose tissue of C57Bl/6 mice fed a HFD for 1, 4, 8, 12 or 16 weeks. C: Gene expression of *Hilpda, Cd68, Adgre1, Itgax, Ccl2* in gonadal adipose tissue of C57Bl/6 mice fed a HFD for 20 weeks (LFD, n=8; HFD, n=10). D: Immunohistochemical staining of HILPDA in gonadal adipose tissue from C57Bl/6 mice fed a HFD for 20 weeks. Right panel is without primary antibody. Bar graphs are presented as mean ± SEM. Gene expression levels in LFD mice were set to one. *P < 0.05, **P ≤ 0.001, ***P≤ 0.0001. LFD: low fat diet, HFD: high fat diet.

### Hilpda is responsive to lipids in macrophages

To further investigate the regulation of HILPDA by fatty acid and intralipid loading, we treated RAW 264.7 macrophages and primary peritoneal macrophages with fatty acids. Confirming the transcriptomics data, oleate markedly upregulated *Hilpda* mRNA in RAW 264.7 and peritoneal macrophages (Fig 2A). Similarly, intralipid treatment significantly induced *Hilpda* mRNA in both types of macrophages (Fig 2B). The induction of HILPDA by intralipid in RAW264.7 and peritoneal macrophages was particularly evident at the protein level (Fig 2C). Since *Hilpda* was originally identified as hypoxia-induced gene 2 (*Hig2*)^15^, and hypoxic areas are characteristic of obese adipose tissue^16^, we tested the effect of intralipid loading in combination with chemical hypoxia. As shown in figure 2D, chemical hypoxia and intralipid synergistically increased *Hilpda* mRNA.

**Figure 2.**
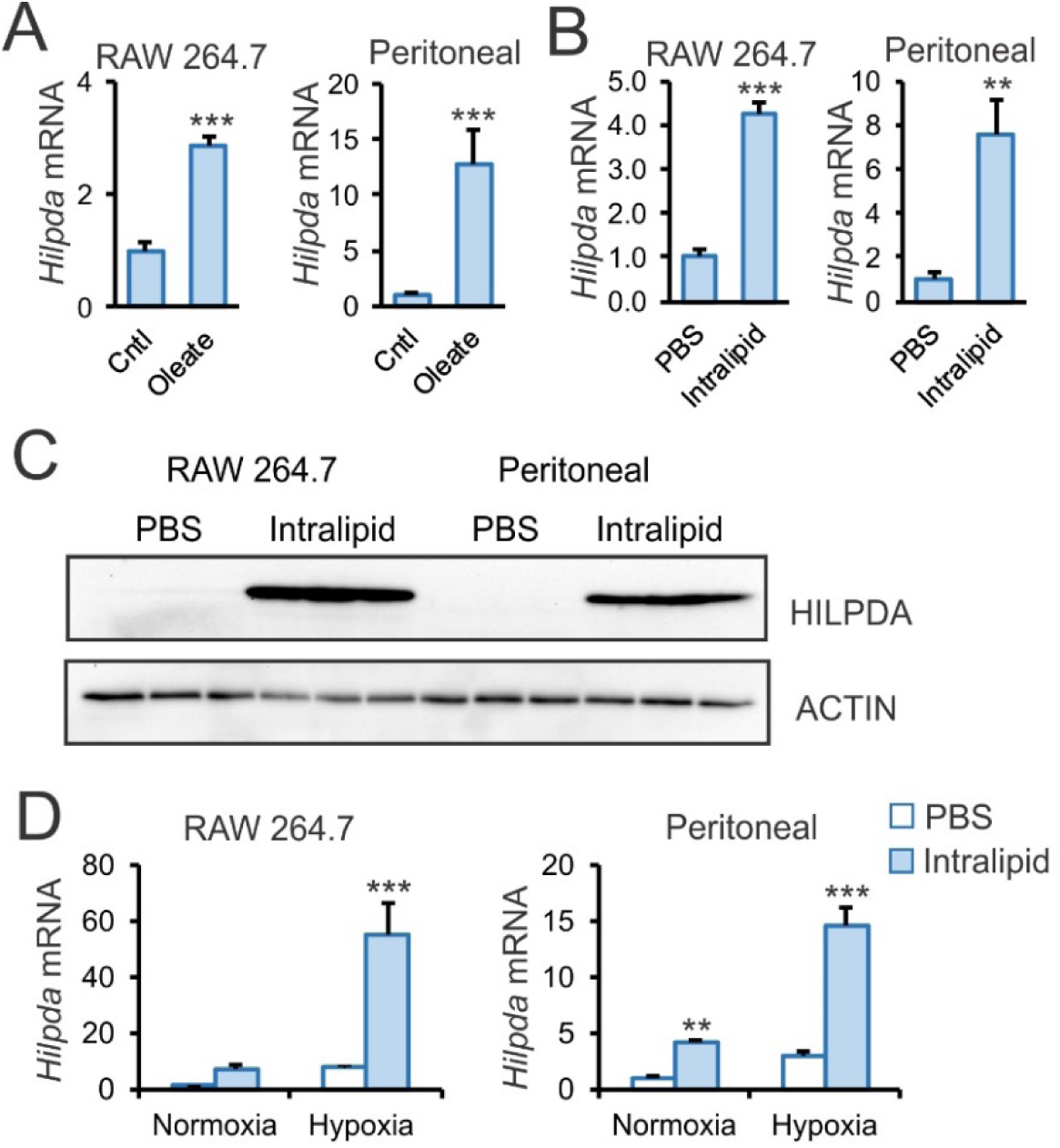
*Hilpda* is responsive to lipids in macrophages. *Hilpda* mRNA expression in RAW 264.7 and C57Bl/6 mouse peritoneal macrophages exposed to 250μM oleate (A) or 1mM intralipid (B) for 6h versus BSA control (cntl) or PBS. HILPDA protein expression in RAW 264.7 and peritoneal macrophages exposed to 1mM intralipid or PBS for 6h. ACTIN was used as loading control. D: *Hilpda* mRNA expression in RAW 264.7 and C57Bl/6 mouse peritoneal macrophages exposed to 1mM intralipid or PBS in combination with normoxia or chemical hypoxia induced by 100μM 2,2’-bipyridyl for 6h. Bar graphs are presented as mean ± SD. Gene expression levels in control (cntl), PBS treatments or PBS with normoxia were set to one. The effect of treatment was significant in D. **P ≤ 0.001, ***P≤ 0.0001.

### Macrophage specific Hilpda deficiency impairs lipid droplet accumulation

We next switched to bone-marrow derived macrophages (BMDMs) as a robust primary *in vitro* model, as they enable studying the effects of *Hilpda* deficiency. Similar to RAW 264.7 and peritoneal macrophages, BMDMs from wild-type C57Bl/6 mice showed a strong synergistic upregulation of *Hilpda* mRNA after intralipid loading and chemical hypoxia (Fig 3A). To be able to study the effects of *Hilpda* deficiency in macrophages, we generated mice with a myeloid-specific *Hilpda* inactivation (*Hilpda*^ΔMΦ^) by crossing *Hilpda*^flox/flox^ with mice expressing Cre-recombinase driven by the LysM promoter. BMDMs obtained from *Hilpda*^ΔMΦ^ mice and their *Hilpda*^flox/flox^ littermates were lipid loaded with a combination of oleate and palmitate for 12 hours to induce maximal lipid droplet formation. The Cre-mediated excision in BMDMs led to an approximate 80% reduction in *Hilpda* mRNA (Fig 3B) and a corresponding decrease in HILPDA protein (Fig 3C). Strikingly, staining of neutral lipids by Bodipy in BMDMs showed that lipid droplets were much less visible in fatty acid-loaded *Hilpda*^ΔMΦ^ than *Hilpda*^flox/flox^ macrophages (Fig 3D). Further quantitative analysis indicated that both the number of lipid droplets per cells and the size of the lipid droplets were significantly lower in *Hilpda*^ΔMΦ^ than in *Hilpda*^flox/flox^ macrophages (Fig 3E). The marked reduction of lipid droplets by *Hilpda* deficiency was confirmed using oil red O staining (Fig 3F). In addition, biochemical analysis indicated that triglyceride levels were markedly decreased in fatty acid-loaded *Hilpda*^ΔMΦ^ macrophages compared to *Hilpda*^flox/flox^ (Fig 3G), which was further supported by thin layer chromatography (Sup fig 1A). Taken together, these data show that *Hilpda* deficiency in macrophages leads to a pronounced decrease in lipid storage in lipid droplets.

**Figure 3.**
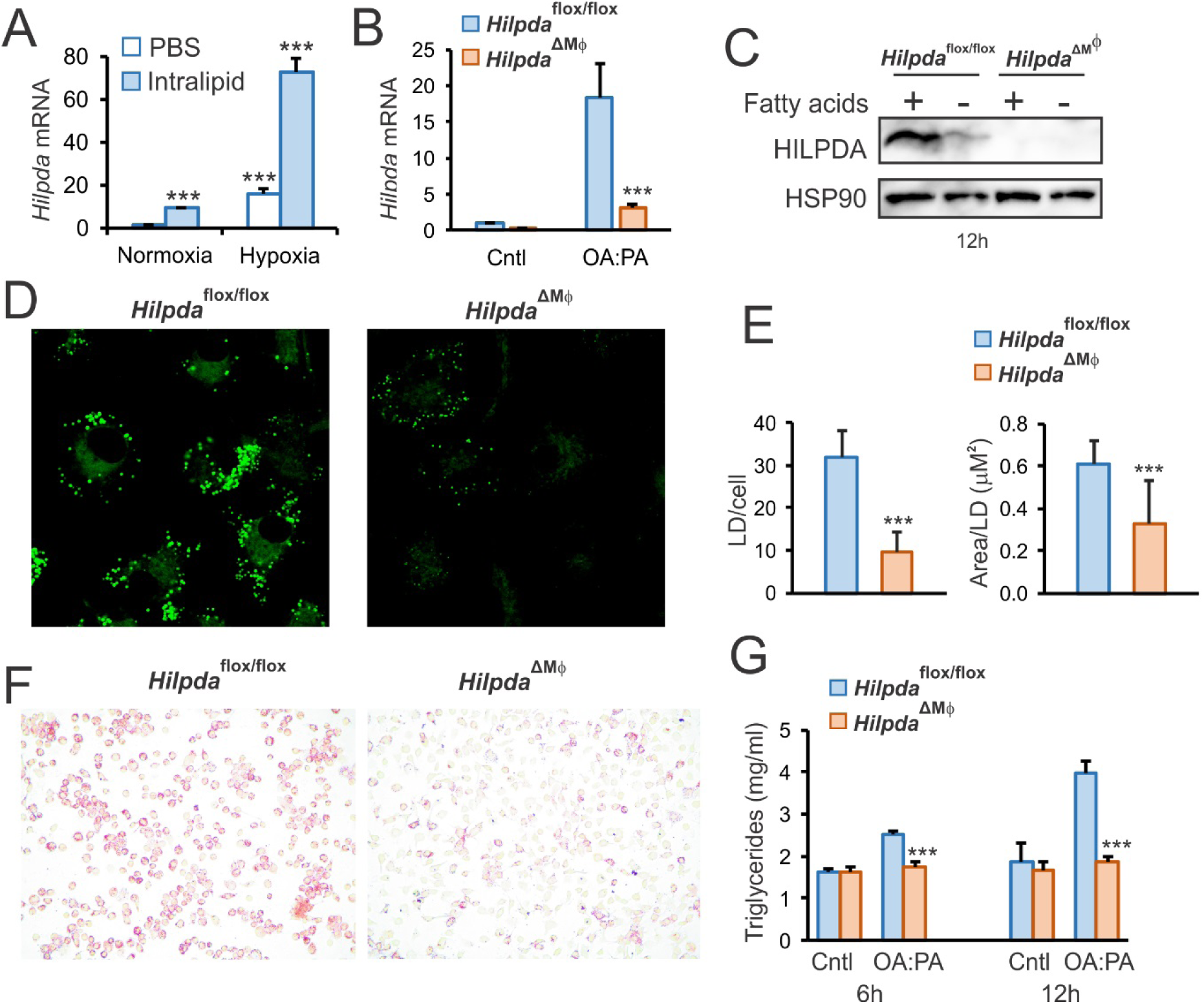
Myeloid-specific *Hilpda* deficiency impairs lipid droplet formation. A: *Hilpda* mRNA expression in C57Bl/6 BMDMs exposed to 1mM intralipid or PBS in combination with normoxia or chemical hypoxia induced by 100μM 2,2’-bipyridyl for 6h. Gene expression of *Hilpda* (B) and protein expression (C) of HILPDA in *Hilpda*^flox/flox^ and *Hilpda*^ΔMΦ^ BMDMs lipid loaded with a mixture of 400μM oleate and 200μM palmitate (oleate:palmitate) or BSA control (cntl) for 12h. Gene expression levels in BSA-treated *Hilpda*^flox/flox^ BMDMs (cntl) are set to one. HSP90 was used as loading control. BODIPY (D and E) and Oil Red O (F) staining in *Hilpda*^flox/flox^ and *Hilpda*^ΔMΦ^ BMDMs lipid loaded with oleate:palmitate or BSA control (cntl) for 24h. G: Triglyceride measurement in BMDMs from *Hilpda*^flox/flox^ and *Hilpda*^ΔMΦ^ lipid loaded with oleate:palmitate or BSA control (cntl) for 6h or 12h. Bar graphs are presented as mean ± SD. Gene expression levels in control (cntl), PBS treatments or PBS with normoxia were set to one. The effect of treatment was significant in B and G. *P < 0.05, ***P≤ 0.0001. OA:PA: oleate:palmitate, LD: lipid droplet.

### HILPDA does not regulate fatty acid uptake or triglyceride synthesis

We next explored potential mechanisms underlying the decrease in lipid storage in *Hilpda*-deficient macrophages. To determine if the reduction in lipid storage in *Hilpda*-deficient macrophages may be due to a decrease in lipid uptake, we measured fatty acid uptake about 30 min after addition of a mixture of oleate and BODIPY-labeled C12 (BODIPY FL). Confocal image analysis showed no difference in fluorescence intensity between *Hilpda*^ΔMΦ^ and *Hilpda*^flox/flox^ macrophages, indicating that *Hilpda* deficiency did not influence fatty acid uptake (Fig 4A, B). In addition, the early induction of gene expression by fatty acids was not different between *Hilpda*^ΔMΦ^ and *Hilpda*^flox/flox^ macrophages, regardless of whether the fatty acids were presented to the cells as free fatty acids (Fig 4C), or intralipid (Sup fig 2), lending further credence to the notion that HILPDA does not regulate fatty acid uptake. Based on these data, combined with the decrease in triglyceride levels in *Hilpda*^ΔMΦ^ macrophages, we hypothesized that HILPDA may have two—not necessarily mutually exclusive—functions: 1) HILPDA may function as an activator of triglyceride synthesis, and/or 2) HILPDA may function as an inhibitor of triglyceride lipolysis. If HILPDA acts by activating triglyceride synthesis in macrophages, suggested before by Maier et al.^17^, it would be expected that *Hilpda* deficiency leads to accumulation of intermediates in the triglyceride synthesis pathways. To explore that possibility, we performed shotgun lipidomics on *Hilpda*^ΔMΦ^ and *Hilpda*^flox/flox^ BMDMs loaded with oleate:palmitate for 24h. Partial least squares discriminant analysis (PLS) clearly separated the two genotypes (Fig 4D), indicating that the lipidomics profiles of the *Hilpda*^ΔMΦ^ and *Hilpda*^flox/flox^ macrophages are very distinct. Interestingly, volcano plot analysis indicated that while a number of lipids were increased in *Hilpda*^ΔMΦ^ macrophages, the majority of lipids were in fact reduced in the *Hilpda*^ΔMΦ^ macrophages (Fig 4E). We reasoned that if HILPDA activates triglyceride synthesis, specific lipid intermediates in the triglyceride synthesis pathway would be expected to accumulate in *Hilpda*^ΔMΦ^ BMDMs. However, phosphatidic acid, diacylglycerols, and triglycerides were significantly downregulated in *Hilpda*^ΔMΦ^ versus *Hilpda*^flox/flox^ macrophages (Fig 4F), as were cholesteryl-esters, while lysophosphatidic acids were hardly detectable. The decrease in triglycerides and diacylglycerol covered the major subspecies within the lipid class (Fig 4G). Taken together, these data suggest that HILPDA probably does not regulate the triglyceride synthesis pathway.

**Figure 4.**
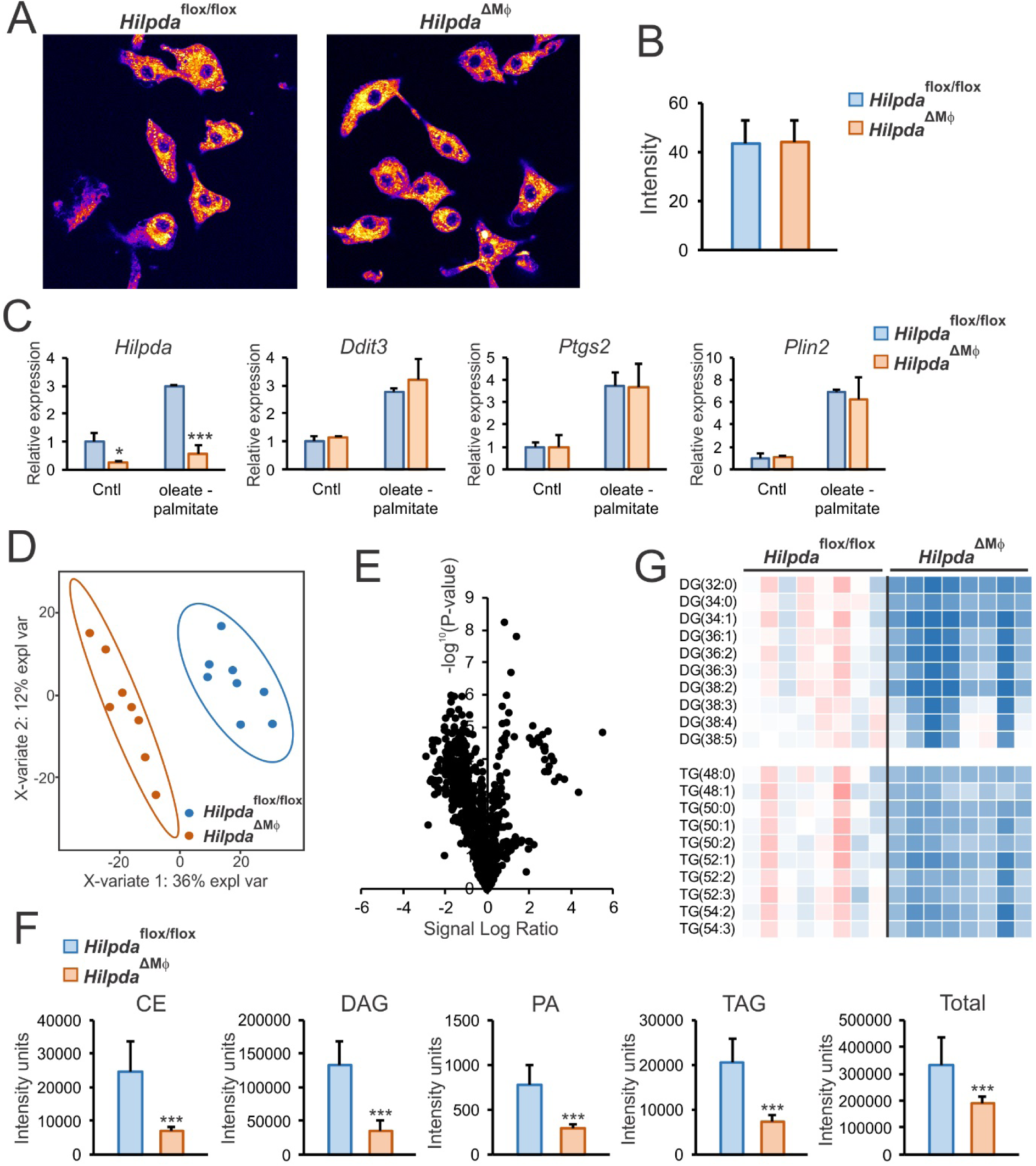
HILPDA does not regulate fatty acid uptake or triglyceride synthesis. A and B: Fatty acid uptake of BODIPY FL in *Hilpda*^flox/flox^ and *Hilpda*^ΔMΦ^ BMDMs after 35 minutes. C: Gene expression of *Hilpda, Ddit3, Ptgs2* and *Plin2* in BMDMs from *Hilpda*^flox/flox^ and *Hilpda*^ΔMΦ^ lipid loaded with oleate:palmitate or BSA control (cntl) for 6h. Partial least square discriminant analysis (D), volcano plot analysis (E), differences in abundancy of CE, DAG, PA, TAG and total lipid species (F) and heatmap of most abundant DG and TG species (G) based on shotgun lipidomics on *Hilpda*^flox/flox^ and *Hilpda*^ΔMΦ^ BMDMs loaded with oleate:palmitate for 24h. Bar graphs are presented as mean ± SD. The effect of treatment was significant in C. *P < 0.05, ***P≤ 0.0001.

### HILPDA regulates lipid droplet mobilization through ATGL inhibition

To further investigate the molecular basis for the decrease in lipid storage in *Hilpda*-deficient macrophages, we determined the trafficking of lipids after loading with a mixture of oleate and BODIPY FL for either 5 or 24 hours. Strikingly, after lipid loading *Hilpda*^flox/flox^ macrophages for 5 hours, the BODIPY FL had accumulated largely in lipid droplets, whereas in *Hilpda*^ΔMΦ^ macrophages, the BODIPY FL was mainly distributed throughout the ER and showed only minor presence in lipid droplet-like structures (Fig 5A). After lipid loading for 24 hours, the size and number of lipid droplets had further increased in *Hilpda*^flox/flox^ macrophages, whereas in *Hilpda*^ΔMΦ^ macrophages, the lipid droplet-like structures that had initially formed at 5h were no longer visible (Fig 5A). These data indicate that *Hilpda*^ΔMΦ^ BMDMs, while being able to take up similar amounts of fatty acids compared to *Hilpda*^flox/flox^ BMDMs, are unable to retain them in lipid droplets. Accordingly, *Hilpda* deficiency in macrophages seems to lead to unstable lipid droplets.

**Figure 5.**
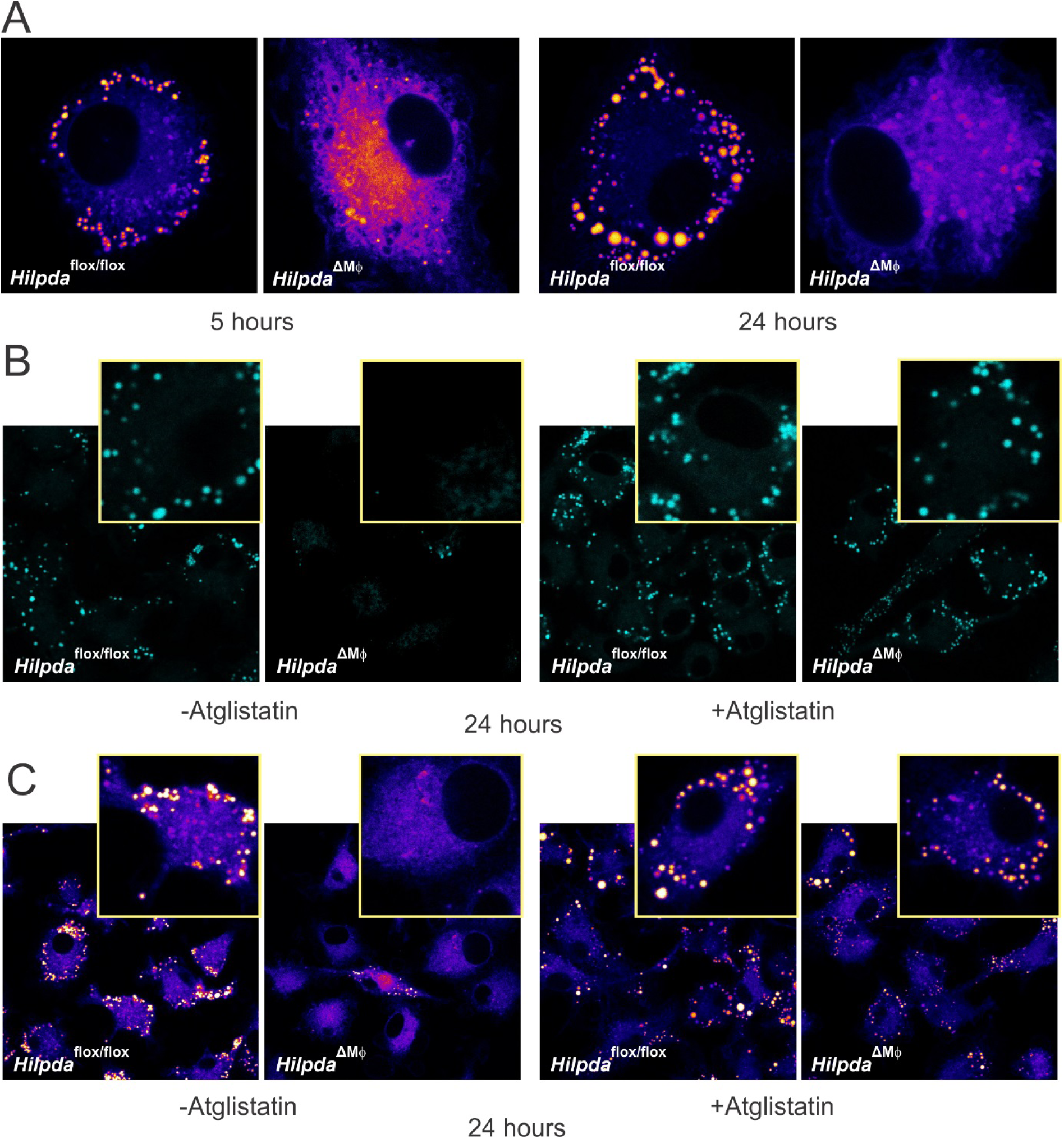
HILPDA regulates lipid droplet mobilization through ATGL inhibition. A: Fatty acid trafficking in *Hilpda*^flox/flox^ and *Hilpda*^ΔMΦ^ BMDMs lipid loaded with 400μM oleate and 20μM BODIPY FL for 5 or 24h. B: BODIPY staining in *Hilpda*^flox/flox^ and *Hilpda*^ΔMΦ^ BMDMs lipid loaded with oleate:palmitate for 24h, treated with 20μM Atglistatin or vehicle. C: Fatty acid trafficking in *Hilpda*^flox/flox^ and *Hilpda*^ΔMΦ^ BMDMs lipid loaded with 400μM oleate and 20μM BODIPY FL for 24h, treated with 20μM Atglistatin or vehicle. LD: lipid droplet.

Using various biochemical and cellular assays, we previously found that HILPDA is able to inhibit ATGL, the rate-limiting enzyme for lipolysis^14^. However, it is still unclear whether HILPDA serves as a physiological regulator of ATGL. To investigate if the decrease in lipid droplets and triglyceride accumulation in *Hilpda*^ΔMΦ^ macrophages is due to enhanced ATGL-mediated lipolysis, we loaded *Hilpda*^ΔMΦ^ and *Hilpda*^flox/flox^ BMDMs with oleate:palmitate in the presence of Atglistatin, a small-molecule inhibitor of ATGL^18^. Strikingly, inhibiting ATGL markedly increased lipid droplets in *Hilpda*^ΔMΦ^ BMDMs (Fig 5B), almost completely rescuing the *Hilpda*^ΔMΦ^ phenotype. Quantitative analysis showed that the LD surface area was not significantly affected by Atglistatin in *Hilpda*^flox/flox^ BMDMs and markedly induced by Atglistatin in *Hilpda*^ΔMΦ^ BMDMs (Sup fig 3A). Similarly, the defective retention of BODIPY FL in lipid droplets in *Hilpda*^ΔMΦ^ macrophages was almost completely abolished by Atglistatin (Fig 5C, Sup fig 3B). The results of these studies suggest that the decrease in lipid droplet and triglyceride accumulation in *Hilpda*^ΔMΦ^ macrophages is caused by accelerated lipid droplet breakdown via enhanced ATGL-mediated lipolysis. Our data thus suggest that HILPDA functions as a potent endogenous inhibitor of ATGL in macrophages.

### HILPDA deficiency promotes respiration

It could be expected that if lipid droplet accumulation in BMDMs is reduced due to enhanced lipolysis, the free fatty acid levels in the cell may rise, thereby stimulating fatty acid dependent gene regulation. Consistent with this notion, the expression of fatty acid-inducible *Gdf15, Cpt1a and Il7r* was significantly higher in lipid-loaded *Hilpda*^ΔMΦ^ than *Hilpda*^flox/flox^ BMDMs (Fig 6A). To analyze this further, we performed transcriptomics on *Hilpda*^ΔMΦ^ and *Hilpda*^flox/flox^ BMDMs loaded with oleate:palmitate for 24h and co-analyzed the data together with a transcriptomics dataset of Robblee et al.^19^ on wild-type BMDMs loaded with stearate for 20h, as well as with the transcriptomics dataset of adipose tissue macrophages isolated from obese mice versus lean mice. In line with enhanced fatty acid-dependent gene regulation in *Hilpda*-deficient macrophages, genes that were highly upregulated after stearate in wild-type BMDMs, such as *Gdf15, Il7r* and *Ddit3*, were also higher in fatty acid loaded *Hilpda*^ΔMΦ^ compared to *Hilpda*^flox/flox^ BMDMs (Fig 6B). In addition, pro-inflammatory genes such as *Ccl2* and *Il1b* were downregulated by stearate in wild-type BMDMs and were also lower in fatty acid loaded *Hilpda*^ΔMΦ^ compared to *Hilpda*^flox/flox^ BMDMs. Interestingly, genes induced by obesity in adipose tissue macrophages, such as *Lpl, Lipa*, and *Itgax*, were only weakly regulated by stearate and by *Hilpda* deficiency, suggesting distinct regulatory mechanisms (Fig 6B).

**Figure 6.**
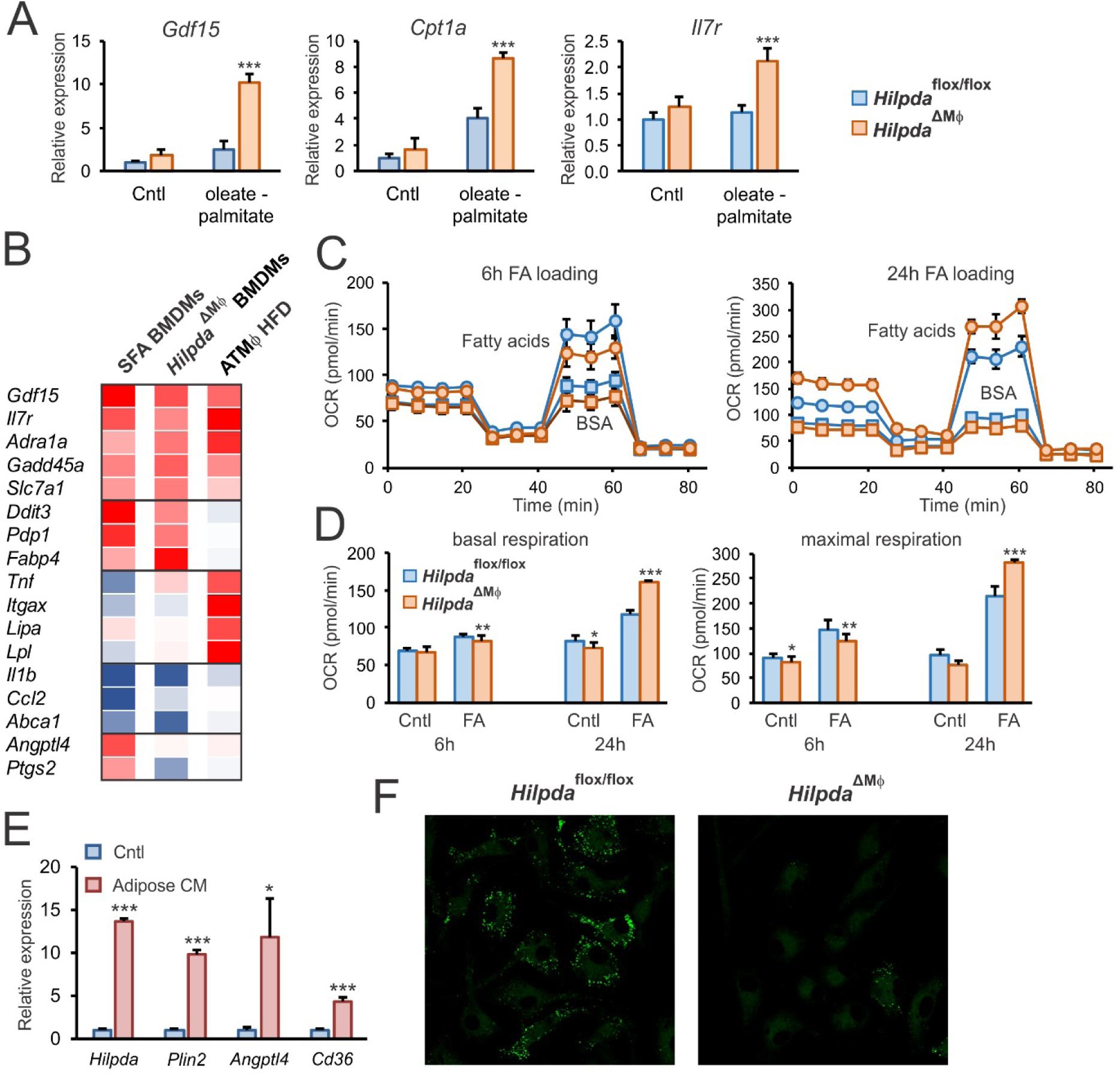
Lack of ATGL inhibition by HILPDA does not affect lipid-induced inflammation, but increases oxidative respiration. A: Gene expression of *Gdf15, Cpt1a, Il7r* and *Fabp4* in *Hilpda*^flox/flox^ and *Hilpda*^ΔMΦ^ BMDMs lipid loaded with oleate:palmitate or BSA control (cntl) for 12h. *Hilpda*^flox/flox^ cntl samples set at one. B: Microarray-based gene expression of relevant genes in WT C57Bl/6 mouse BMDMs loaded with stearate (250μM) for 20h, *Hilpda*^ΔMΦ^ BMDMs loaded with oleate:palmitate for 24h and adipose tissue macrophages isolated from mice fed a HFD vs LFD. Oxygen consumption rate of *Hilpda*^flox/flox^ and *Hilpda*^ΔMΦ^ lipid loaded with oleate:palmitate or BSA control (cntl) for 6h or 24h measured with extracellular flux (C) and corresponding basal and maximal respiration levels (D). Gene expression levels of *Hilpda, Plin2, Angptl4* and *Cd36* in C57Bl/6 mouse BMDMs loaded with adipose conditioned medium or control medium (cntl) for 6h (E) and BODIPY staining after loading with adipose conditioned medium for 24h (F). Untreated controls (cntl) are set to one. Bar graphs are presented as mean ± SD. The effect of treatment was significant in A and D. *P < 0.05, **P ≤ 0.001, ***P≤ 0.0001. FA: fatty acid loading with oleate:palmitate.

Of the 6600 genes that passed the expression threshold, only 49 were induced more than 2 fold in fatty acid loaded *Hilpda*^ΔMΦ^ compared to *Hilpda*^flox/flox^ BMDMs. The apparent limited effect of impaired triglyceride retention in *Hilpda*^ΔMΦ^ BMDMs suggests that the excess fatty acids may be disposed of, for instance by enhanced oxidation. To explore that option, cellular respiration was determined in fatty acid loaded *Hilpda*^ΔMΦ^ and *Hilpda*^flox/flox^ BMDMs. As a marker for oxidative phosphorylation, oxygen consumption of fatty acid loaded *Hilpda*^ΔMΦ^ and *Hilpda*^flox/flox^ BMDMs was measured by extracellular flux analysis during a mitochondrial stress test (Fig 6C). After 6 hours of oleate:palmitate loading, basal respiration in *Hilpda*^ΔMΦ^ and *Hilpda*^flox/flox^ BMDMs was similar to the untreated controls (Fig 6D). Fatty acid loading increased maximal respiration, which was slightly lower in *Hilpda*^ΔMΦ^ and *Hilpda*^flox/flox^ BMDMs. Remarkably, however, after 24h fatty acid loading, basal and maximal respiration were significantly higher in *Hilpda*^ΔMΦ^ compared to *Hilpda*^flox/flox^ BMDMs (Fig 6D), indicating an increased maximal oxidative capacity. These data suggest that the enhanced lipolysis in *Hilpda*^ΔMΦ^ BMDMs is accompanied by increased fatty acid oxidation through oxidative phosphorylation, most likely to limit cellular lipotoxicity.

Our next step was to test whether *Hilpda* might influence lipid droplet accumulation in the context of adipose tissue. To mimic the adipose environment *in vitro*, BMDMs were treated with conditioned medium of adipose tissue explants. Adipose conditioned medium markedly increased *Hilpda* expression, along with the expression of several other lipid sensitive genes, such as *Plin2, Cd36* and *Angptl4* (Fig 6E). Consistent with our previous studies with fatty acid-loaded macrophages, *Hilpda*^ΔMΦ^ BMDMs incubated with adipose conditioned medium showed substantially reduced BODIPY staining compared with *Hilpda*^flox/flox^ BMDMs (Fig 6F). These data suggest that HILPDA may also influence lipid droplet accumulation in macrophages in the context of adipose tissue.

### Myeloid-specific deficiency of Hilpda decreases lipid droplets in ATMs without altering adipose tissue inflammation

To enable studying the effect on *Hilpda* deficiency in macrophages in vivo, we used *Hilpda*^ΔMΦ^ mice and their *Hilpda*^flox/flox^ littermates. As expected, myeloid-specific inactivation of *Hilpda* led to a significant decrease in *Hilpda* expression in the stromal vascular fraction of adipose tissue, but not in the adipocyte fraction (Fig 7A). To test the functional consequences of macrophage *Hilpda* deficiency in the context of obesity-induced adipose tissue inflammation and foam cell formation, *Hilpda*^ΔMΦ^ mice and their *Hilpda*^flox/flox^ littermates were rendered obese and insulin resistant by high fat feeding for 20 weeks, using a low fat diet as control. Bodyweight gain (Fig 7B), feed intake (Fig 7C), and liver and adipose tissue weights (Fig 7D) were not different between *Hilpda*^ΔMΦ^ and *Hilpda*^flox/flox^ littermates. Consistent with the data shown in Fig 1C, high fat feeding increased *Hilpda* mRNA in adipose tissue. Interestingly, the relative increase in *Hilpda* mRNA was considerably lower in *Hilpda*^ΔMΦ^ adipose tissue than in *Hilpda*^flox/flox^ adipose tissue (Fig 7E), suggesting that the increase in *Hilpda* expression by high fat feeding is mainly driven by its expression in macrophages. Immunoblot for HILPDA confirmed this notion by showing markedly reduced HILPDA protein levels in *Hilpda*^ΔMΦ^ versus *Hilpda*^flox/flox^ adipose tissue (Fig 7F).

**Figure 7.**
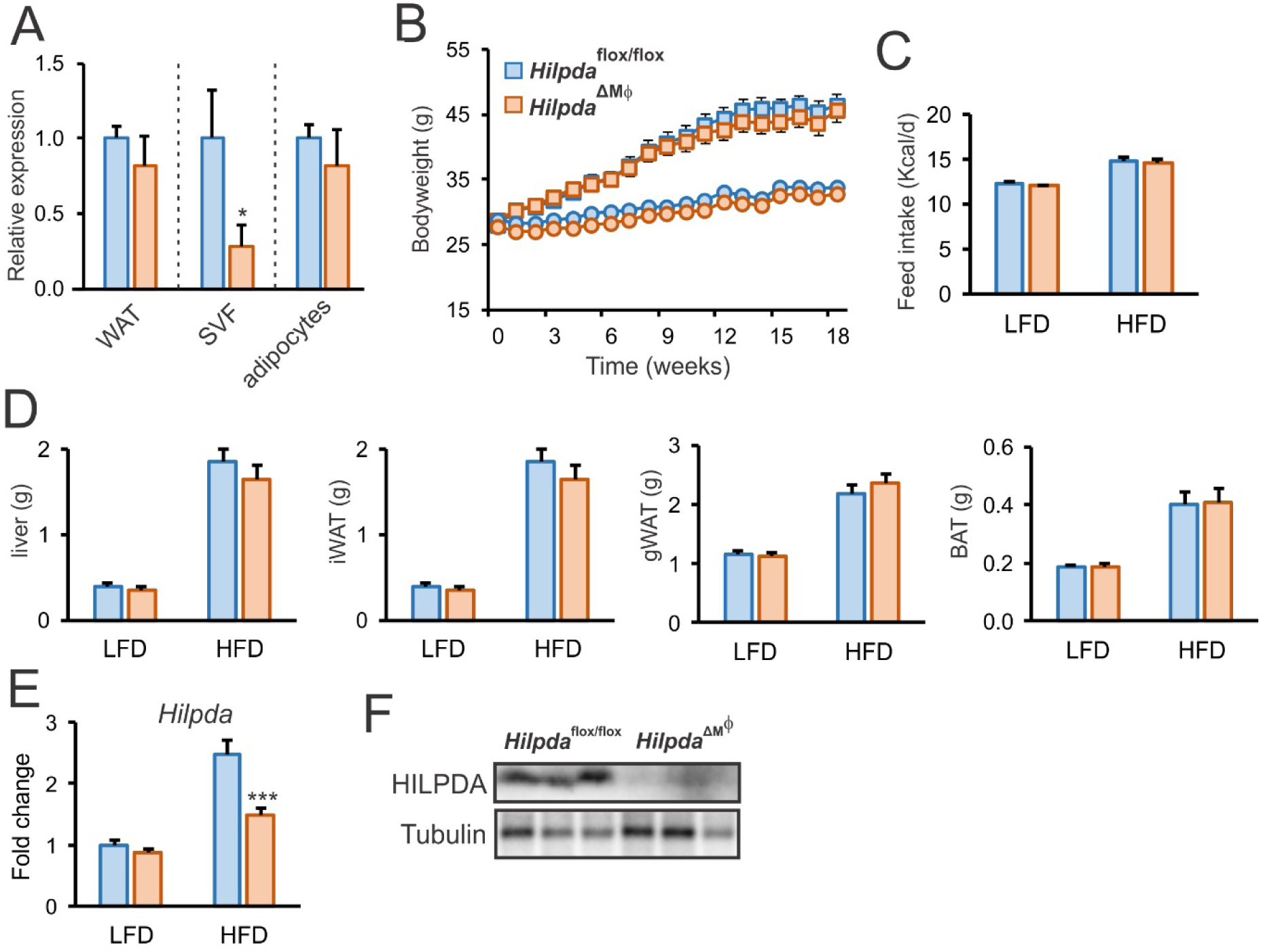
*Hilpda* deficiency in myeloid cells results in a decrease in HILPDA in gWAT after high fat feeding, without influencing body and organ weight. A: Relative gene expression of *Hilpda* in whole gWAT and corresponding stromal vascular fraction and adipocyte fraction of *Hilpda*^flox/flox^ and *Hilpda*^ΔMΦ^ fed normal chow. Levels of expression in *Hilpda*^flox/flox^ for every fraction set to one. Body weight (B), feed intake (C) and weight of liver, iWAT, gWAT and BAT (D) in *Hilpda*^flox/flox^ and *Hilpda*^ΔMΦ^ mice fed a LFD or HFD for 20 weeks. *Hilpda* gene expression in gWAT of *Hilpda*^flox/flox^ and *Hilpda*^ΔMΦ^ mice fed a LFD or HFD for 20 weeks (E) and HILPDA protein expression in gWAT of *Hilpda*^flox/flox^ and *Hilpda*^ΔMΦ^ mice fed a HFD for 20 weeks (F). Bar graphs are presented as mean ± SEM. The effect of diet was significant in C, D and E. *P < 0.05, ***P≤ 0.0001. WAT: white adipose tissue, SVF: stromal vascular fraction, LFD: low fat diet, HFD: high fat diet, iWAT: inguinal adipose tissue, gWAT: gonadal adipose tissue, BAT: brown adipose tissue.

Based on the studies in BMDMs, we hypothesized that lipid accumulation would be reduced in adipose tissue macrophages from *Hilpda*^ΔMΦ^ mice compared to *Hilpda*^flox/flox^ mice. Indeed, Oil Red O staining showed that lipid droplet content was significantly lower in adipose tissue macrophages isolated from HFD-fed *Hilpda*^ΔMΦ^ mice compared with HFD-fed *Hilpda*^flox/flox^ mice (Fig 8A,B). Interestingly, however, the decrease in lipid droplets was not associated with any change in the secretion of the classical inflammatory cytokines IL6 and TNFα (Fig 8C). These data indicate that *Hilpda* deficiency reduces lipid accumulation in adipose tissue macrophages but does not have any effect on their *ex vivo* inflammatory properties.

**Figure 8.**
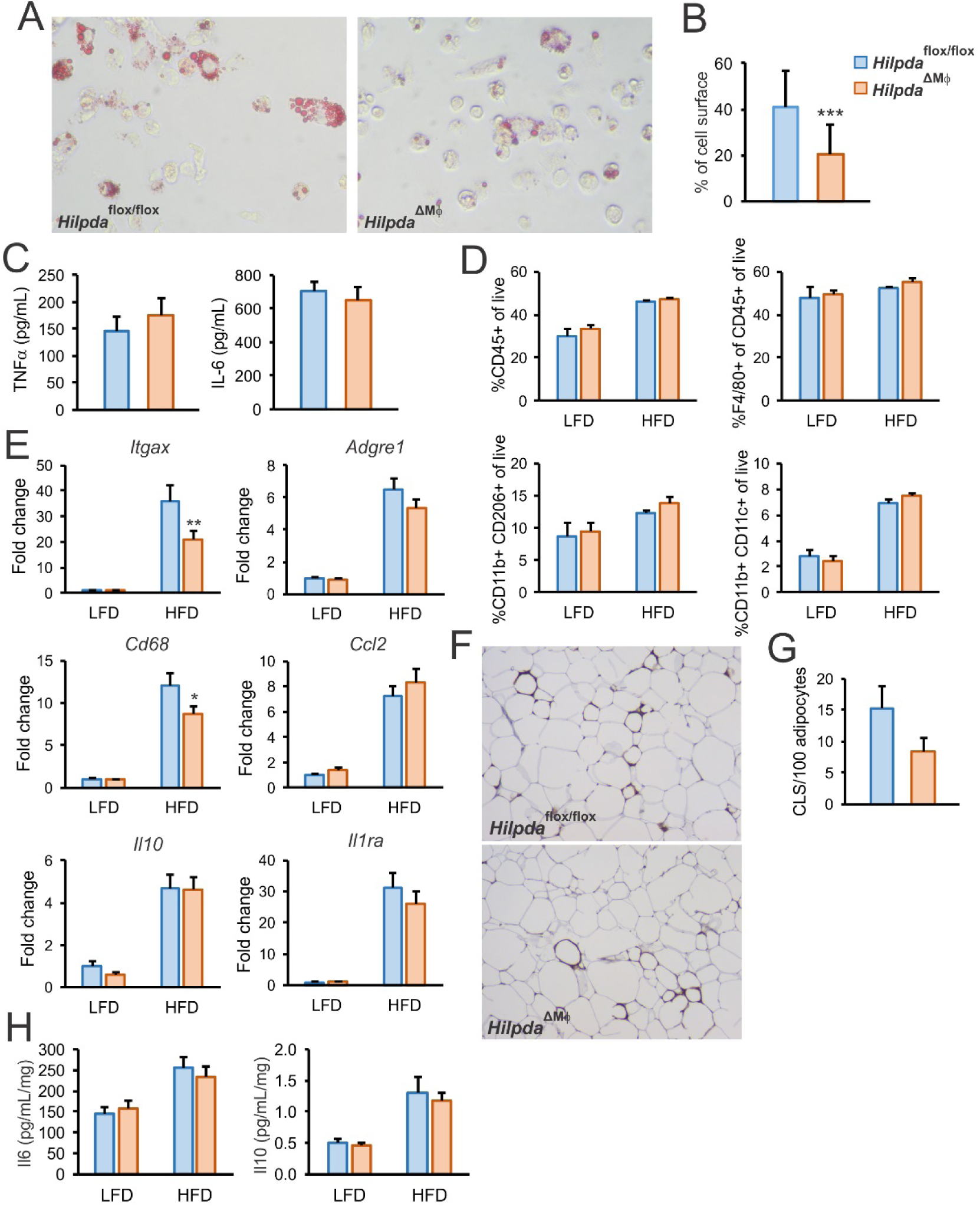
Myeloid-specific deficiency of *Hilpda* decreases lipid droplets in ATMs without altering adipose tissue inflammation. Oil red O staining of adipose tissue macrophages isolated from *Hilpda*^flox/flox^ and *Hilpda*^ΔMΦ^ mice fed a HFD for 20 weeks (A and B). Data are mean ± SD. C: Corrected TNFα and IL-6 secretion of adipose tissue macrophages isolated from *Hilpda*^flox/flox^ and *Hilpda*^ΔMΦ^ mice fed a HFD for 20 weeks. D: Flow cytometry based percentages of CD45+, CD45+F4/80+, CD11b+CD206+ and CD11b+CD11c+ population in the SVF of gWAT from *Hilpda*^flox/flox^ and *Hilpda*^ΔMΦ^ mice fed a LFD or HFD for 20 weeks. Gene expression of *Itgax, Adgre1, Cd68, Ccl2, Il10, Il1ra* (E), density of CLSs in gWAT coupes stained for F4/80 (F and G, only for HFD) and secretion of IL-6 and IL-10 (H) in gWAT explants of *Hilpda*^flox/flox^ and *Hilpda*^ΔMΦ^ mice fed a LFD or HFD for 20 weeks. Expression in gWAT of the *Hilpda*^flox/flox^ LFD group is set to one, bar graphs are presented as mean ± SEM. The effect of diet was significant in D, E and H. *P < 0.05, **P≤ 0.001, ***P≤ 0.0001. LFD: low fat diet, HFD: high fat diet, CLS: crown-like structure.

To investigate the potential impact of macrophage HILPDA on adipose tissue inflammation *in vivo*, we performed flow cytometry analysis of the stromal vascular fraction isolated from the adipose tissue of the various groups of mice. The results showed an increased percentage of populations of CD45+, CD11b+CD206+ and CD11b+CD11c+ cells by high fat feeding, but no clear differences in the percentages of these populations between *Hilpda*^ΔMΦ^ and *Hilpda*^flox/flox^ mice (Fig 8D). To further examine the influence of macrophage *Hilpda* deficiency on the inflammatory status of adipose tissue, the expression of selected genes was determined in adipose tissue of *Hilpda*^ΔMΦ^ and *Hilpda*^flox/flox^ mice. Interestingly, the expression of both inflammatory macrophage marker *Itgax* (Cd11c) and general macrophage marker *Cd68* was significantly lower in adipose tissue of *Hilpda*^ΔMΦ^ mice versus *Hilpda*^flox/flox^ mice fed a HFD, while the general macrophage marker *Adgre1* (F4/80) showed a trend towards a decreased expression (Fig 8E). Despite being induced by high fat feeding, adipose expression of other genes involved in pro- or anti-inflammatory signalling, such as *Gdf15, Il10, Arg1, Ccl2* and *Il1ra* was not different between *Hilpda*^ΔMΦ^ and *Hilpda*^flox/flox^ mice (Fig 8E, Sup fig 4). Expression of *Adipoq* and *Leptin* also was not different between *Hilpda*^ΔMΦ^ and *Hilpda*^flox/flox^ mice (Sup fig 4).

To further investigate the inflammatory status of adipose tissue, the density of crown-like structures was determined in adipose tissue of *Hilpda*^ΔMΦ^ and *Hilpda*^flox/flox^ mice fed a HFD. A trend towards lower density was found in the *Hilpda*^ΔMΦ^ mice (Fig 8F), which, however, did not reach statistical significance (Fig 8G). As a last measurement of inflammatory status, we collected adipose tissue explants and measured the *ex vivo* release of cytokines. Despite the fact that high fat feeding significantly stimulated the release of IL10 and IL6 by, no difference in IL10 and IL6 release could be observed between adipose tissue explants derived from *Hilpda*^ΔMΦ^ and *Hilpda*^flox/flox^ mice (Fig 8H).

Finally, to determine whether macrophage *Hilpda* deficiency has any influence on obesity-induced metabolic derailments, we measured plasma metabolic parameters and assessed glucose tolerance in *Hilpda*^ΔMΦ^ and *Hilpda*^flox/flox^ mice fed the low and high fat diets. High fat feeding significantly increased plasma levels of cholesterol, triglycerides, non-esterified fatty acids, glucose, insulin, and leptin (Fig 9A). However, no difference in these parameters were observed between *Hilpda*^ΔMΦ^ and *Hilpda*^flox/flox^ mice, either on a low or high fat diet (Fig 9A). Similarly, although high fat feeding caused a marked decrease in glucose tolerance, no differences were observed between *Hilpda*^ΔMΦ^ and *Hilpda*^flox/flox^ mice (Fig 9B).

**Figure 9.**
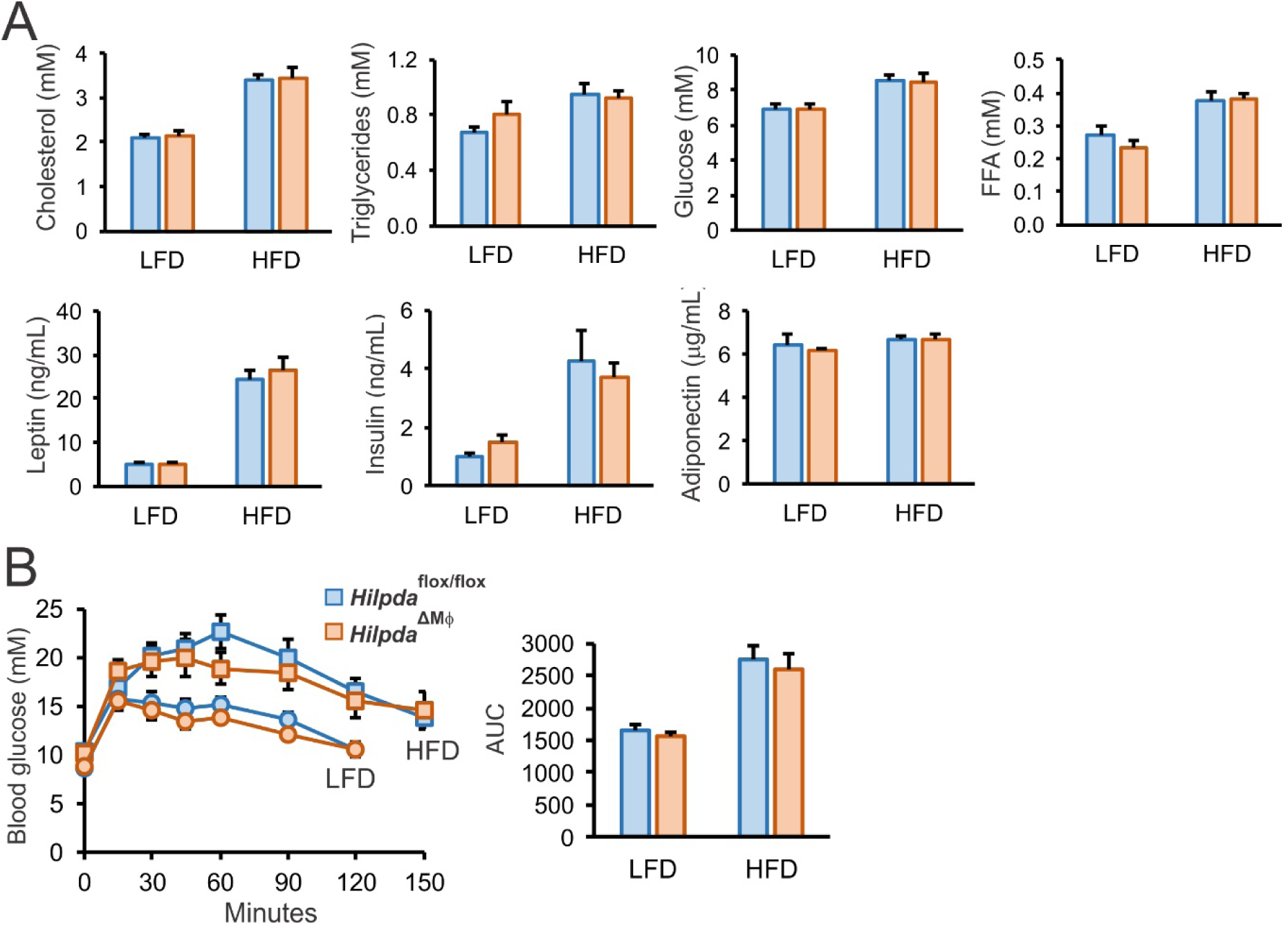
*Hilpda* deficiency in myeloid cells does not influence glucose tolerance after high fat feeding. A: Plasma levels of cholesterol, triglycerides, glucose, free-fatty acids, leptin, insulin and adiponectin in *Hilpda*^flox/flox^ and *Hilpda*^ΔMΦ^ mice fed a LFD or HFD for 20 weeks. B: Intra-peritoneal glucose tolerance test adiponectin in *Hilpda*^flox/flox^ and *Hilpda*^ΔMΦ^ mice after 18 weeks of either LFD or HFD feeding. Bar graphs are presented as mean ± SEM. The effect of diet was significant in cholesterol, triglycerides, glucose, FFA, leptin and insulin in A and AUCs in B. LFD: low fat diet, HFD: high fat diet, FFA: free fatty acids.

Taken together, our data indicate that myeloid-specific *Hilpda* deficiency reduces lipid accumulation in adipose tissue macrophages. Along with reduced adipose foam cell formation, *Hilpda* deficiency may cause a slight decrease in the macrophage population in adipose tissue. However, *Hilpda* deficiency does not influence the inflammatory status of adipose tissue, nor does it have any effect on obesity-induced metabolic complications.

## DISCUSSION

This study was aimed at determining the role of HILPDA in lipid accumulation in (adipose tissue) macrophages and explore the potential causal relationship between macrophage lipid accumulation and the development of adipose tissue inflammation and insulin resistance during obesity. Here, we provide evidence that *Hilpda* deficiency in macrophages disrupts stable lipid droplet formation after lipid loading. We further show that the marked decrease in lipid storage in *Hilpda*-deficient macrophages is due to impaired retention of lipids in the lipid droplets. Specifically, we find that the decrease in lipid storage in *Hilpda*-deficient macrophages can be almost completely abolished by inhibition of ATGL, demonstrating that HILPDA is an endogenous and physiological inhibitor of ATGL in macrophages. The lack of inhibition of ATGL-mediated lipolysis in *Hilpda*-deficient macrophages in turn leads to increased oxidative phosphorylation. Strikingly, despite reducing lipid storage in adipose tissue macrophages, *Hilpda* deficiency in macrophages does not alter the inflammatory status of adipose tissue in diet-induced obesity, arguing against the notion that lipid accumulation in adipose tissue macrophages promotes adipose tissue inflammation and associated insulin resistance.

During obesity, macrophages infiltrate the adipose tissue and take up adipocyte-released lipids. Consequently, lipid laden adipose tissue macrophage, or adipose foam cells, are a distinct macrophage population found in crown-like structures in murine and human obesity^7,8,20^. This macrophage subpopulation shows a characteristic activation that has been suggested to be mediated by the uptake and accumulation of excess lipids released by adipocytes, which likely represents an attempt to dispose of the lipid excess^21–23^. The use of mice deficient in *Hilpda* in macrophages allowed us to investigate to what extent lipid accumulation in adipose tissue macrophages may drive adipose tissue inflammation and associated insulin resistance. Since lipid droplet formation likely serves as a cytoprotective mechanism to prevent lipotoxic effects caused by lipid intermediates or free fatty acids^24^, it could be hypothesized that the enhanced lipid droplet breakdown in *Hilpda*-deficient macrophages results in elevated inflammation. However, *Hilpda* deficiency not only reduced intracellular triglyceride levels but also the levels of potentially lipotoxic intermediates, such as DAG, that may contribute to reducing the inflammatory responses of the macrophage. This reduction is most likely achieved via enhanced fatty acid oxidation in the macrophage, a metabolic pathway that in itself is also known to impact the inflammatory status of macrophages^25^. Accordingly, it could also be reasoned that *Hilpda* deficiency should lead to reduced inflammation via these pathways. Intriguingly, however, no effect of *Hilpda* deficiency was observed on cytokine release by adipose tissue macrophages, on the percentage of different macrophage populations in adipose tissue, on inflammatory gene expression in adipose tissue, and on cytokine release by adipose tissue explants. Also, genes typically elevated in adipose tissue macrophages from obese mice, such as *Lipa, Lpl*, and *Itgax*, were not induced by fatty acid loading or altered upon *Hilpda* deficiency. The data argue against the notion that excessive lipid accumulation in adipose tissue macrophages is the major driver of adipose tissue inflammation and of the composition of macrophage populations in obese adipose tissue. Rather, the unique profile of adipose tissue macrophages may be determined by other factors active in the obese adipose tissue environment, the identity of which requires further study.

As indicated above, *Hilpda*-deficient macrophages not only had a reduced number and size of lipid droplets, but also exhibited a marked decrease in intracellular levels of all of the major lipid species, including triglycerides, diacylglycerol, phosphatidic acid, and cholesteryl-esters. Since fatty acid uptake was unaltered by *Hilpda* deficiency, these data suggest that the oxidation of lipid is enhanced in *Hilpda*-deficient macrophages. Indeed, we find increased oxidative respiration in *Hilpda*-deficient macrophages. Previously, a strong link was made between ATGL activity and fatty acid oxidation, at least in liver and heart. Specifically, it was found that ATGL-mediated lipolysis activates a transcriptional network involving PGC-1α/PPAR-α that controls fatty acid oxidation and mitochondrial biogenesis^26–29^. Accordingly, it is likely that the loss of ATGL inhibition is directly responsible for the enhanced oxidative capacity, reducing the total intracellular lipid load. In general, increased lipolysis as well as increased oxidative respiration are two traits essential for macrophage polarization towards alternative, M2-like phenotypes, which has been suggested to be protective in the context of adipose tissue inflammation^30–32^. Interestingly, in our experiments, increased oxidative respiration seemed a mere consequence following overactive ATGL-mediated lipolysis, and did not contribute to any anti-inflammatory effects in the context of adipose tissue inflammation. Although increased oxidation of fatty acids is often proposed as an alternative cytoprotective mechanism in lipid-laden macrophages, the complex interplay between fatty acid oxidation, concomitant ROS formation, and ER stress has not yet led to clear evidence for such an effect^25^. In practice, the combination of possible pro-inflammatory effects of dysfunctional triglyceride storage by overactive ATGL-mediated lipolysis and the possible anti-inflammatory effect of enhanced fatty acid oxidation might cancel out each other and explain the absence of any clear inflammatory changes in *Hilpda*-deficient macrophages in obese adipose tissue.

Apart from fatty acid oxidation, ATGL has also been linked to the autophagic degradation of lipid droplets, termed lipophagy^33^. It was suggested that ATGL acts as a signaling node to promote lipophagy, which then controls bulk lipid droplet breakdown. Whether HILPDA, via ATGL, connects to lipophagy requires further study.

In contrast to the cytoprotective effect of normal lipid droplet formation, the adverse effects of excessive triglyceride storage becomes apparent in ATGL-/-macrophages, underlining the importance of functional ATGL in macrophages. ATGL-/-macrophages cannot effectively break down triglyceride stores and show substantial triglyceride accumulation, leading to mitochondrial dysfunction and apoptosis, endoplasmic reticulum (ER) stress, reduced macrophage migration, and decreased phagocytosis ability^34–37^. Macrophages deficient in the ATGL activator CGI-58 (a.k.a. ABHD5) also have elevated lipid storage and decreased phagocytic capacity, yet show no signs of mitochondrial apoptosis and ER stress, suggesting that TG accumulation per se does not drive mitochondrial dysfunction^38^. Our data show that *Hilpda* deficiency, despite leading to markedly reduced lipid storage, raises markers of ER stress, suggesting that triglyceride storage protects against lipid-induced ER stress. Presumably, the mechanism leading to ER stress is different in *Hilpda*-deficient macrophages as compared with ATGL/CGI-58-deficient macrophages.

HILPDA was initially identified in a subtractive hybridization screen for hypoxia-induced genes in human cervical cancer cells and was later found to be associated with lipid droplets^9,39^. We identified *Hilpda* as a novel PPARa target gene in liver^40^. In addition, *Hilpda* is well expressed in adipocytes^10,12^. Several studies have shown that overexpression of *Hilpda* increases intracellular lipid storage in cells^9,10,40^. In the present study, *Hilpda* emerged from a screen for genes elevated by obesity in adipose tissue macrophages and upregulated in macrophages by fatty acid treatment. The induction of *Hilpda* by fatty acids and the subsequent inhibition of triglyceride hydrolysis is likely part of an effort of the cell to effectively store excess energy and neutralize the potentially reactive free fatty acids and thus a crucial component of the lipid buffering capacity of macrophages. Of note, the minor effect of Atglistatin on lipid storage in wildtype BMDMs suggest that ATGL is almost fully inhibited in macrophages, showing the importance of controlling triglyceride hydrolysis.

We previously found that HILPDA is able to inhibit ATGL in biochemical assays, analogous to the ATGL inhibitor G0/G1 switch gene 2 (G0S2), with which ATGL shares extensive sequence homology^14,41^. However, the inhibitory action of HILPDA was low compared to G0S2, which raised questions on the physiological relevance of HILPDA as an inhibitor of ATGL. Our studies demonstrate that HILPDA acts as a potent endogenous inhibitor of ATGL-mediated lipolysis in macrophages. Our preliminary data also suggest that the expression level of *Hilpda* in BMDMs is at least 30-fold higher than the expression of *G0s2*. A number of questions emerge from this work. First, why does HILPDA, despite allegedly being a much weaker ATGL inhibitor than G0S2, have such a marked influence on lipid storage in macrophages? We hypothesize that HILPDA may require an interaction with an auxiliary factor for full activity. Further research is necessary to identify the mechanism for the differential potency of HILPDA in cell-free systems compared to live cells. Second, what is the reason for having two related ATGL inhibitors? It seems that at least in certain cells, such as hepatocytes, HILPDA and G0S2 co-exist. Inasmuch as HILPDA and G0S2 are induced by different stimuli, they are likely active under different circumstances. So far there is no evidence for any functional dependency between the two proteins. In addition, using FRET-FLIM analysis, we were unable to demonstrate any physical interaction between G0S2 and HILPDA (our unpublished data). Further research is necessary to better characterize the relationship and relative roles of these two homologous proteins in different cell types.

There is evidence that lipid droplets may influence the immunological properties of macrophages^42^. Interestingly, activation of macrophages by immunological stimuli such as TLR ligands enhances fatty acid uptake and lipid droplet formation, coupled with a decrease in triglyceride lipolysis and an increase in fatty acid synthesis^43^. Newly synthesized fatty acids have been suggested to play a role in the proinflammatory activation and response of macrophages^44^. According to our preliminary data, expression of *Hilpda* in macrophage is markedly induced by immunological stimuli. Induction of *Hilpda* by immunological stimuli may promote lipid storage and be part of a mechanism to regulate macrophage function via modulation of lipid droplet homeostasis. Further research should investigate the role of HILPDA in lipid droplet homeostasis and pathogenic macrophage activation.

In conclusion, our data demonstrate that HILPDA is a lipid-induced physiological inhibitor of ATGL-mediated lipolysis in macrophages. In obese mice, HILPDA uncouples lipid storage in adipose tissue macrophages from inflammation and metabolic dysregulation. Overall, our data question the importance of lipid storage in adipose tissue macrophages in obesity-induced inflammation and metabolic dysregulation.

## METHODS

### Animal studies

Animal studies were performed using purebred wild-type C57BL/6 animals (Jackson Laboratories, Bar Harbor, ME), *Hilpda*^ΔMΦ^ mice and their *Hilpda*^flox/flox^ littermates. *Hilpda*^flox/flox^ were acquired (Jackson Laboratories, Bar Harbor, ME; *Hilpda*^*tm*^*1.1*^*Nat*^, #017360) and crossed with C57Bl/6 mice for at least 5 generations. Thereafter, the *Hilpda*^flox/flox^ were crossed with *lysM*-Cre transgenic mice (Jackson Laboratories, Bar Harbor, ME; B6.129P2-Lyz2tm1(cre)Ifo/J, #004781) to generate mice with a mature myeloid cell-specific Cre-mediated deletion of *Hilpda*. Mice were individually housed under normal light-dark cycles in temperature- and humidity-controlled specific pathogen-free conditions. Mice had ad libitum access to food and water.

Male *Hilpda*^ΔMΦ^ mice aged 9-12 weeks and their male *Hilpda*^flox/flox^ littermates were placed on a high fat diet for 20 weeks to induce obesity and insulin resistance. From earlier studies it is known that fasting glucose values of mice fed a high fat diet differs on average 3mM (± 8 mM – 11mM) compared to mice fed a low fat diet. Differences in responses lead to a standard deviation around 2mM or higher. For the power calculation, we used a one-way ANOVA with a significance level of 0.05 and a power of 90%, leading to an estimation of around *n* = 11 mice needed per group. To allow compensation for unforeseen circumstances or potential loss of mice during the study, *n* = 12 mice were included per group. Therefore, 12 mice per genotype were randomly allocated using an online randomisation tool to either a standardized high fat diet or a low fat diet (formula D12451 and formula D12450H respectively, Research Diets, New Brunswick, USA; γ-irradiated with 10-20 kGy) for 20 weeks.

Body weight and food intake were assessed weekly. At the end of the study, mice were anaesthetised with isoflurane and blood was collected via orbital puncture in tubes containing EDTA (Sarstedt, Nümbrecht, Germany). Subsequently, mice were immediately euthanized by cervical dislocation, after which tissues were excised, weighed and frozen in liquid nitrogen or prepared for histology. Samples from liquid nitrogen were stored at −80°C. All animal experiments were approved by the local animal welfare committee of Wageningen University (AVD104002015236, 2016.W-0093.001). The experimenter was blinded to group assignments during all analyses.

### Intraperitoneal glucose tolerance test

In study 3, an intraperitoneal glucose tolerance test was performed after 18 weeks. Mice were fasted for 5 hours and blood was collected via tail bleeding at 0, 15, 30, 45, 60, 90 and 120 minutes after i.p. injection of 1g/kg bodyweight glucose (Baxter, Deerfield, IL, USA). Blood glucose was measured with a GLUCOFIX Tech glucometer and glucose sensor test strips (GLUCOFIX Tech, Menarini Diagnostics, Valkenswaard, The Netherlands). A time point of 150 minutes after injection of glucose was added for the high fat diet fed groups.

### Plasma measurements

Blood collected in EDTA tubes was spun down for 15 minutes at 5000 RPM at 4°C, plasma was aliquotted and stored in −80°C until measurement of cholesterol (Liquicolor, Human GmbH, Wiesbaden, Germany), triglycerides (Liquicolor), glucose (Liquicolor), NEFAs (NEFA-HR set R1, R2 and standard, WAKO Diagnostics, Instruchemie, Delfzijl, The Netherlands), adiponectin (ELISA duoset kit, R&D Systems, Bio-techne, MN, USA), leptin (ELISA duoset kit, R&D Systems) and insulin (ultra-sensitive mouse insulin ELISA kit, Crystal Chem Inc., IL, USA) following manufacturer’s instructions.

### gWAT explants and isolation of adipose tissue macrophages

For SVF, adipocytes and adipose tissue macrophages isolation, gonadal adipose tissue (gWAT) was collected and kept in with Dulbecco’s modified Eagle’s medium (DMEM, Corning, NY, USA), supplemented with 1% penicillin/streptomycin (p/s, Corning) and 1% FFA-free Bovine Serum Albumin (BSA fraction V, Roche via Merck, Darmstadt, Germany) on ice. gWAT explants were taken into culture for 24h in DMEM, supplemented with 10% fetal calf serum (FCS, BioWest, Nuaillé, France) and 1% p/s. Supernatant was stored for ELISA measurements or as conditioned medium. For high fat diet groups, the stromal vascular fractions were isolated by digesting gWAT for 45 minutes in Roswell Park Memorial Institute (RPMI)-1630 medium (Lonza, Basel, Zwitserland) supplemented with 10% FCS, 1% p/s, 0.5% FFA-free BSA, 1M CaCl2, 1M HEPES and 0.15% collagenase (from *Clostridium histolyticum*, Merck). Per three mice of the same group, gWAT was pooled after digestion, filtered through a 100μm cell strainer and centrifuged at 200*g* for 10 min. Floating mature adipocytes were removed and stored separately and stromal vascular pellet was resuspended in erythrocyte lysis buffer and subsequently washed twice in phosphate buffered saline (PBS, Corning) supplemented with 0.5% FFA-free BSA and 2mM EDTA. Resulting stromal vascular fractions were used to isolate ATMs using mouse anti-F4/80-FITC antibodies (Miltenyi Biotec, Bergisch Gladbach, Germany), anti-FITC MicroBeads (Miltenyi Biotec) and MS columns (Miltenyi Biotec) on an OctoMACS™Cell Separator system (Miltenyi Biotec). ATMs were cultured for 24h in RPMI-1630 supplemented with 10% FCS and 1% P/S. ATMs were either cultured for 2h after which cells were washed with PBS, fixed in 3.7% paraformaldehyde and stained with Oil red O following standard protocols, or were cultured for 24h to obtain supernatants.

### Flow cytometry of SVF

Before isolation of ATMs, SVF pools were resuspended in PBS containing 0.5% BSA and 2mM EDTA and 500 000 cells were sampled and stained with antibodies against CD45-ECD (Beckman Coulter, Brea, CA, USA), F4/80-FITC, CD206-APC, CD11c-PE-Cy7 and CD11b-PE (Biolegend, San Diego, CA, USA). Samples were measured on a flow cytometer (FC500, Beckman Coulter) and results were analyzed using Kaluza analysis software 2.1 (Beckman Coulter).

### Histological studies

Samples of gWAT for histological analysis were fixed in 3.7% paraformaldehyde immediately upon collection, embedded in paraffin, sectioned and stained with hematoxylin eosin according to standard protocols. After preincubation with 20% normal goat serum, paraffin-embedded sections were incubated at 4°C overnight with antibodies for F4/80 (MCA497G, Bio-Rad Laboratories, Hercules, CA, USA), HILPDA (sc-137518 HIG2 Antibody (C-14), Santa-Cruz Biotechnology, Dallas, TX, USA Biotechnology) or CD68 (AbD Serotec, Bio-Rad Laboratories) dissolved in PBS supplemented with 1% BSA (Merck). Anti-rat or anti-rabbit IgG conjugated to HRP (Cell Signaling Technology Danvers, MA, USA) were used as secondary antibody. Negative control were prepared without using primary antibody.

### Isolation and stimulation of peritoneal macrophages and BMDMs

To harvest peritoneal macrophages, 8-12 week old WT C57Bl/6 mice were injected intraperitoneally with 1mL 4% thioglycolic acid. Three days post-injection, mice were anesthetised with isoflurane and euthanized by CO_2_. Peritoneal cells were harvested by washing the peritoneal cavity with ice-cold RPMI-1630 supplemented with 10% heat-inactivated FCS (BioWest) and 1% p/s. Cells were plated after lysis of erythrocytes and non-adherent cells were washed away three hours post plating. To isolate BMDMs, 8-12 week old *Hilpda*^ΔMΦ^ mice and their *Hilpda*^flox/flox^ littermates were euthanized by cervical dislocation. Both femurs and hind legs were isolated at the hip joint, keeping femur and tibia intact. Bone marrow was extracted from the femur and tibia and differentiated in DMEM, supplemented with 10% FCS, 1% p/s and 15% L929 conditioned medium. After seven days of differentiation, BMDMs were scraped and plated as appropriate.

### Cell culture experiments

RAW 264.7 macrophages were cultured in DMEM supplemented with 10% FCS and 1% p/s. Palmitate (Merck) and oleate (Merck) were solubilized using EtOH and KOH and conjugated to FFA-free BSA in sterile water (Versol, Aguettant, Lyon, France) at 37°C for 30 min. Palmitate was used in concentrations of 200, 250 or 500μM. Oleate was used in a concentration of 250μM or 400μM together with 20 μM BODIPY-FL C12 (Thermo Fisher Scientific Scientific, MA, USA) for fatty acid trafficking experiments. A mixture of oleate and palmitate (oleate:palmitate) was made in a ratio of 1:2 and used in a final concentration of 600 μM. Intralipid (Fresenius Kabi AB, Uppsala, Sweden) was used in a concentration of 1 or 2mM. Chemical hypoxia was induced by the addition of 100μM iron chelator 2,2’-bipyridyl (Merck). Atglistatin (Merck) was used in a concentration of 20μM in 100% DMSO and cells were pre-treated for 2 hours before fatty acid loading. 24 hour treatments containing Atglistatin were refreshed every 12 hours. All cells were washed with PBS (Corning) after treatment. BMDMs were stained with Oil Red O following standard procedures.

### Confocal Imaging

To visualise fatty acid uptake, accumulation and trafficking, BMDMs were plated on 8-well μ glass bottom slides (Ibidi, Martinsried, Germany). Confocal imaging was performed on a Leica confocal TCS SP8 X system equipped with a 63× 1.20 NA water-immersion objective lens. Images were acquired using 1,024 × 1,024 pixels with pinhole set at 1 Airy Unit (AU). Excitation of the fluorescent probes used in this study was performed using white light laser (WLL, 50% laser output) selecting the 488 nm laser line. Fluorescence emission was detected using internal Hybrid (HyD) detector selecting a spectral window from either 520 - 580 nm (fatty acid uptake) or from 510 – 565 nm (fatty acid trafficking).

Fatty acid uptake was measured on paraformaldehyde fixed cells after 35 minutes incubation with the QBT™ Fatty acid uptake assay kit (Molecular Devices, California, USA) according to manufacturer’s instructions. Image analysis was performed on Fiji. Pixels were selected for analysis using Otsu threshold, mean intensity was quantified. The WLL laser line (488 nm) was set at a laser power of 0.2%. The pinhole was adjusted at 5.7 AU for fluorescence intensity measurements, whereas confocal imaging was done with a pinhole of 1 AU.

Fatty acid trafficking was assessed after lipid loading for 5h and 24h with 400 μM oleate and 20 μM BODIPY® FL C12, treated either with vehicle or Atglistatin. The WLL laser line (488 nm) was set at a laser power of 1.6% for 5 h incubated cells and 0.3 % for 24 h incubated cells. Cells were washed with PBS, fixed for 15 min with 3.7% formaldehyde and mounted with Vectashield-H (Vector Laboratories, Peterborough, UK). Fire LUT was applied using Fiji (https://fiji.sc/).

To assess fatty acid accumulation, BMDMs treated with oleate:palmitate were washed with PBS and fixed for 15 minutes with 3.7% paraformaldehyde. Fixed cells were stained with 2ug/mL BODIPY® 493/503 (Thermo Fisher Scientific) and mounted with Vectashield-H (Vector Laboratories). Images were processed and analyzed with Fiji. Briefly, images were converted to binary, watershed and LD size and number was measured with particle analysis set 0.07 μm2-infinity.

### Extracellular flux assay

Extracellular flux of lipid-loaded BMDMs was measured using the Agilent Seahorse XF96 Analyzer (Agilent Technologies, Santa Clara, CA, USA). Briefly, cells were seeded in a density of 200 000 cells per well in XF-96 plates (Agilent Technologies), treated appropriately and kept in a 37°C/5% CO_2_ incubator. An hour before the measurement, cells were washed and cultured in Seahorse XF base medium (Agilent Technologies) without sodium bicarbonate, supplemented with 25mM glucose and 2mM L-glutamine for one hour at 37°C in a non-CO_2_ incubator. For the mitochondrial stress test, the following compounds were added during four injections: oligomycin (1.5uM), FCCP (1.5uM), pyruvate (1mM), antimycin A (2.5uM) and rotenone (1.25uM). The OCR was automatically measured by the sensor cartridge at baseline and following injections. Calculations were made using the Seahorse XF-96 software Wave Desktop 2.6 (Agilent Technologies).

### Real-time PCR

For cells, total RNA was isolated using TRIzol® Reagent (Invitrogen, ThermoFisher Scientific). For tissues, total RNA was isolated using the RNeasy Micro Kit (Qiagen, Venlo, The Netherlands). cDNA was synthesized from 500ng RNA using the iScript cDNA kit (Bio-Rad Laboratories, Hercules, CA, USA) according to manufacturer’s instructions. Real time polymerase chain reaction (RT-PCR) was performed with the CFX96 or CFX384 Touch™ Real-Time detection system (Bio-Rad Laboratories), using a SensiMix™ (BioLine, London, UK) protocol for SYBR green reactions. Mouse *36b4* expression was used for normalization.

### Immunoblotting

Cell or tissue protein lysates were separated by electrophoresis on pre-cast 4-15% polyacrylamide gels and transferred onto nitrocellulose membranes using a Trans-Blot® Semi-Dry transfer cell (all purchased from Bio-Rad Laboratories), blocked in non-fat milk and incubated overnight at 4°C with primary antibody for HILPDA (Santa-Cruz Biotechnology), ACTIN (Cell Signaling Technology), TUBULIN (Cell Signaling Technology) or HSP90 (Cell Signaling Technology). Membranes were incubated with secondary antibody (Anti-rabbit IgG, HRP-linked Antibody, 7074, Cell Signaling Technology) and developed using Clarity ECL substrate (Bio-Rad Laboratories). Images were captured with the ChemiDoc MP system (Bio-Rad Laboratories).

### Enzyme-linked immunosorbent assay (ELISA)

DuoSet sandwich ELISA kits for TNFα, IL10 and IL6 (R&D systems) were used to measure cytokine concentrations in cell or explant supernatant according to manufacturer’s instructions. Data was normalized for the amount of adipose tissue macrophages by determining the concentration of DNA per well (Quant-iT dsDNA Assay Kit high sensitivity, Thermo Fisher Scientific) and normalized for gWAT explants to the weight per explant.

### Lipidomics

Lipidomics analysis was performed as described (Herzog, 2016). The HPLC system consisted of an Ultimate 3000 binary HPLC pump, a vacuum degasser, a column temperature controller, and an auto sampler (Thermo Fisher Scientific). The column temperature was maintained at 25°C. The lipid extract was injected onto a “normal phase column” LiChrospher 2×250-mm silica-60 column, 5 μm particle diameter (Merck) and a “reverse phase column” Acquity UPLC HSS T3, 1.8 _m particle diameter (Waters, Milford, MA, USA). A Q Exactive Plus Orbitrap (Thermo Fisher Scientific) mass spectrometer was used in the negative and positive electrospray ionization mode. Nitrogen was used as the nebulizing gas. The spray voltage used was 2500 V, and the capillary temperature was 256 _C. S-lens RF level: 50, auxiliary gas: 11, auxiliary temperature 300°C, sheath gas: 48, sweep cone gas: 2. In both the negative and positive ionization mode, mass spectra of the lipid species were obtained by continuous scanning from m/z 150 to m/z 2000 with a resolution of 280,000 full width at half maximum (FWHM). Data was analyzed and visualised using R programming language (https://www.r-project.org). Heat-maps were created using the package “gplots” and partial least squares regression analysis was performed using the R package “mixOMICS”.

### Microarray analyses

Microarray analysis was performed on a several experiments: 1) Peritoneal macrophages treated with various fatty acids (500 μM) for 6 hours. 2) Peritoneal macrophages treated with intralipid (2mM) for 6 hours. 3) BMDM samples from *Hilpda*^ΔMΦ^ mice and *Hilpda*^flox/flox^ mice lipid loaded with oleate:palmitate (600μM) for 12 and 24 hours. RNA was isolated as described above and purified with the RNeasy Micro kit from Qiagen. Integrity of the RNA was verified with RNA 6000 Nano chips using an Agilent 2100 bioanalyzer (Agilent Technologies). Purified RNA (100 ng per sample) was labeled with the Whole-Transcript Sense Target Assay kit (Affymetrix, Santa Clara, CA, USA; P/N 900652) and hybridized to an Affymetrix Mouse Gene 1.0 arrays or 2.1 ST array plate (Affymetrix). Hybridization, washing, and scanning were carried out on an Affymetrix GeneTitan platform according to the manufacturer’s instructions. Normalization of the arrays was performed with the Robust Multi-array Average method^45,46^. Probe sets were redefined according to Dai et al.^47^ based on annotations provided by the Entrez Gene database. Array data have been submitted to the Gene Expression Omnibus (accession numbers pending).

A publicly available dataset (GSE77104) was downloaded from Gene Expression Omnibus and further processed as described above to obtain individual gene expression data. The microarray analysis of the adipose tissue macrophages (GSE84000) is already described elsewhere (Boutens, 2016).

Data for the 3D scatterplot of signal log ratio’s (Fig 1A) was created using R programming language (https://www.r-project.org) and the R package “plot3D”.

### Statistical analysis

Data are represented as means ±SD or SEM as indicated. Statistical analyses were carried out using an unpaired Student’s *t* test or two-way ANOVA followed by Bonferroni’s post hoc multiple comparisons test, if genotype and diet or genotype and treatment both were found significant (GraphPad Software, San Diego, CA, USA). A value of p < 0.05 was considered statistically significant.

## Acknowledgements

We would like to thank Shohreh Keshtkar, Jenny Jansen, Anneke Hijmans and Jacqueline Ratter for their technical assistance.

## Supplemental data

**Sup fig1.**
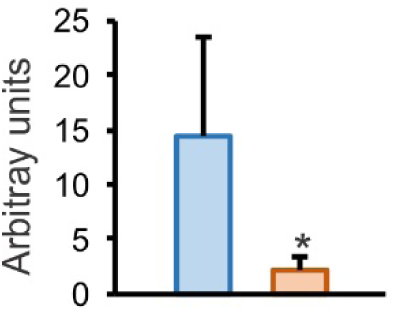
A: Quantification of triglycerides on Thin Layer Chromatography plates in *Hilpda*^flox/flox^ and *Hilpda*^ΔMΦ^ BMDMs lipid loaded with oleate:palmitate for 24h. B: BODIPY staining in *Hilpda*^flox/flox^ and *Hilpda*^ΔMΦ^ BMDMs treated with palmitate for 6h or 24h. Bar graphs are presented as mean ± SD. *P < 0.05.

**Sup fig2.**
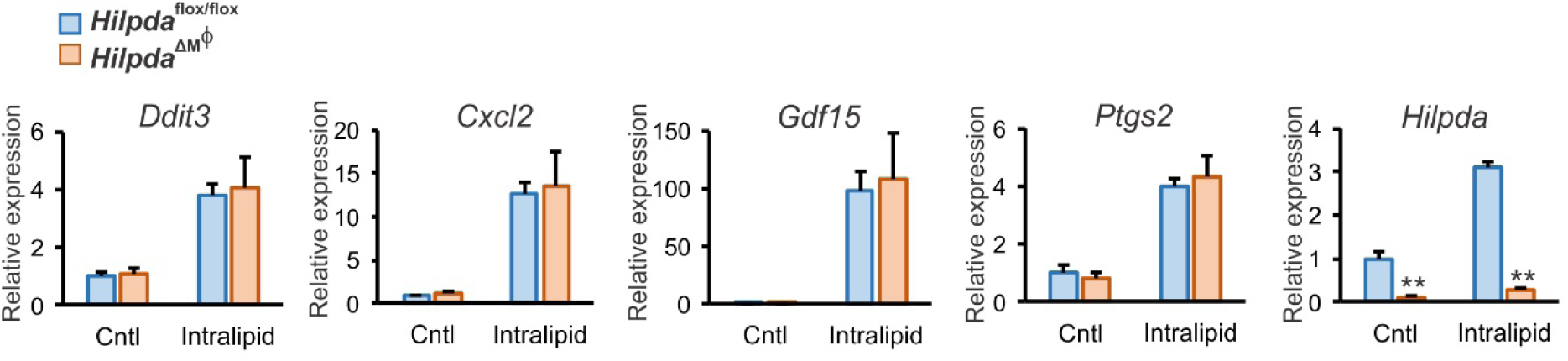
A: Gene expression of *Ddit3, Cxcl2, Gdf15, Ptgs2* and *Hilpda* in *Hilpda*^flox/flox^ and *Hilpda*^ΔMΦ^ BMDMs treated with 1mM intralipid or PBS control (Cntl) for 6h. Bar graphs are presented as mean ± SD. The effect of treatment was significant. **P ≤ 0.001

**Sup fig3.**
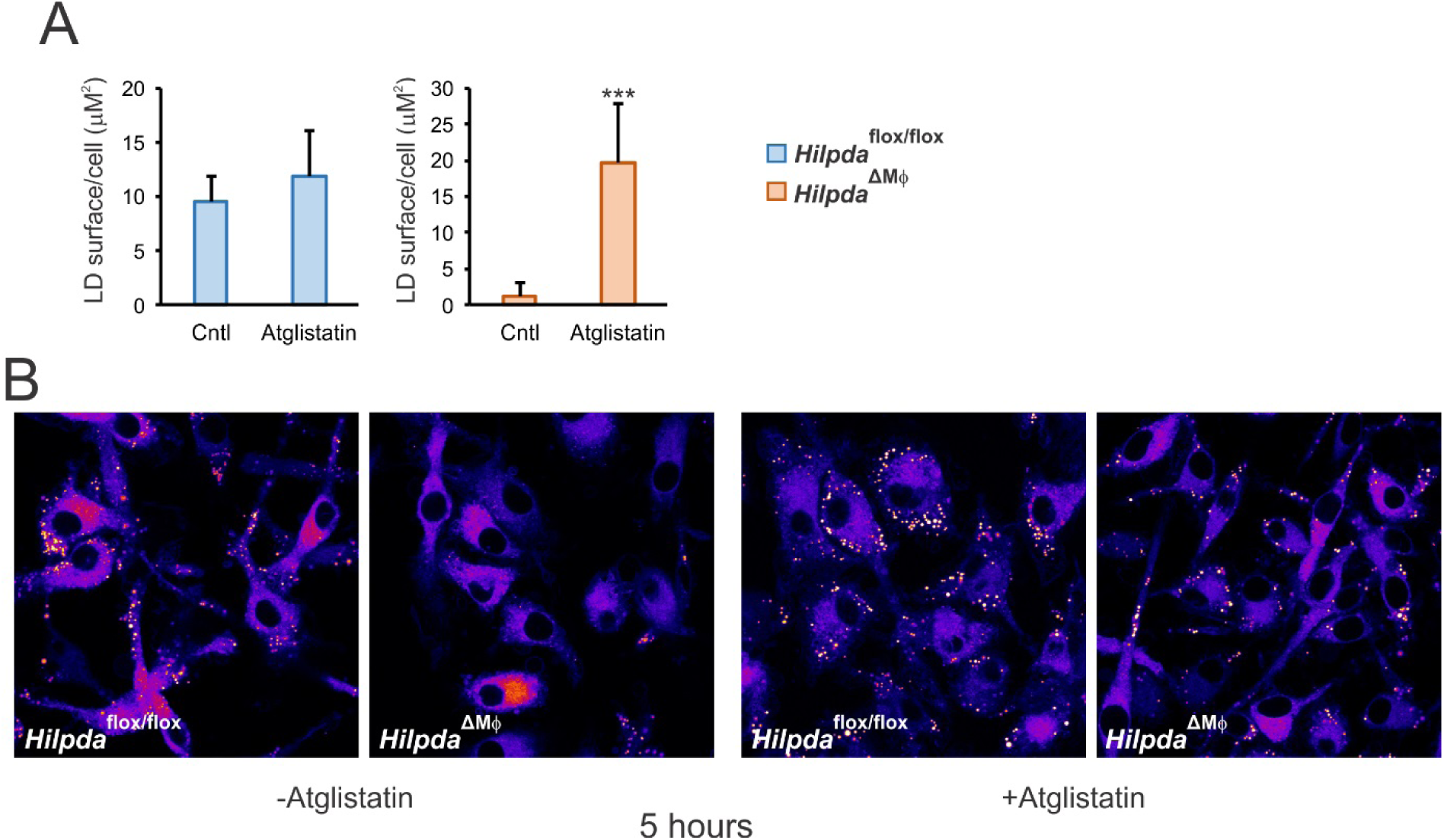
A: Quantification of the amount of lipid droplet surface area per cell in *Hilpda*^flox/flox^ and *Hilpda*^ΔMΦ^ BMDMs lipid loaded with oleate:palmitate for 24h and treated with 20μM atglistatin or vehicle (DMSO). B: BODIPY FL trafficking and incorporation in lipid droplets in *Hilpda*^flox/flox^ and *Hilpda*^ΔMΦ^ BMDMs lipid loaded with 400μM oleate and 20μM BODIPY FL, treated with 20μM atglistatin or vehicle (DMSO) for 5 hours. Bar graphs are presented as mean ± SD.

**Sup fig4.**
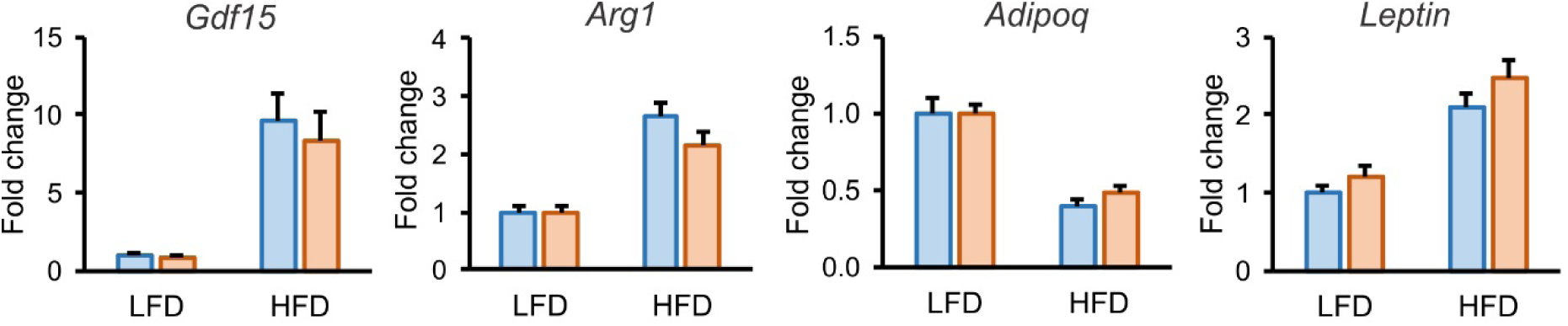
A: Gene expression of *Gdf15, Arg1, Adipoq* and *Leptin* in gWAT of *Hilpda*^flox/flox^ and *Hilpda*^ΔMΦ^ mice fed a LFD or HFD for 20 weeks. Gene expression levels in gWAT from *Hilpda*^flox/flox^ fed a LFD diet are set to one. Bar graphs are presented as mean ± SEM. The effect of diet was significant.

